# A Universal Free-Degree Orientation Extrusion Head Enables Conformal and Non-Planar Bio-Additive Manufacturing toward Adaptive and Future-Ready Bioprinting

**DOI:** 10.64898/2026.06.23.734010

**Authors:** Gopinathan Janarthanan, Rashik Chand, Sanjairaj Vijayavenkataraman

## Abstract

Conventional extrusion-based 3D bioprinting encounters limitations in fabricating intricate tissue architectures due to fixed nozzle diameters and fixed deposition orientations. These constraints restrict conformal printing on curved or non-planar surfaces and often necessitate support-intensive fabrication strategies. This work introduces a mechanically simplified extrusion platform inspired by the swivel jet nozzle, featuring a free-degree-of-orientation extrusion head termed the universal extrusion head (Univ-Ex head), coupled with a modular nozzle architecture. The Univ-Ex head employs a swivel-like mechanical design that enables orientation freedom without external actuation in its current implementation, thereby minimizing mechanical complexity while supporting deposition on physiologically relevant, non-planar geometries. Multiple nozzle concepts were developed through comparative CAD iterations, with two representative geometries—a flat nozzle and a conical nozzle—selected for experimental validation. The platform is evaluated through parametric CAD design, stereolithography-printed prototypes, proof-of-concept extrusion experiments, and fluid dynamics simulations performed using FLOW-3D software. Numerical and experimental results demonstrate stable filament formation and clear diameter-dependent extrusion behavior, while simulations further confirm the feasibility of angled and non-planar deposition. A variable-diameter nozzle concept is proposed as a forward design direction to enable real-time adjustment of bioink flow rate and deposition resolution in principle; however, the present study intentionally validates the system using fixed-diameter nozzle variants to maintain stable numerical and experimental boundary conditions. A gear-integrated Univ-Ex head is also presented as a forward upgrade and demonstrated as a single-piece prototype. Collectively, this work establishes a scalable, hardware-focused pathway toward conformal bio-additive manufacturing.

**Graphical Abstract:** 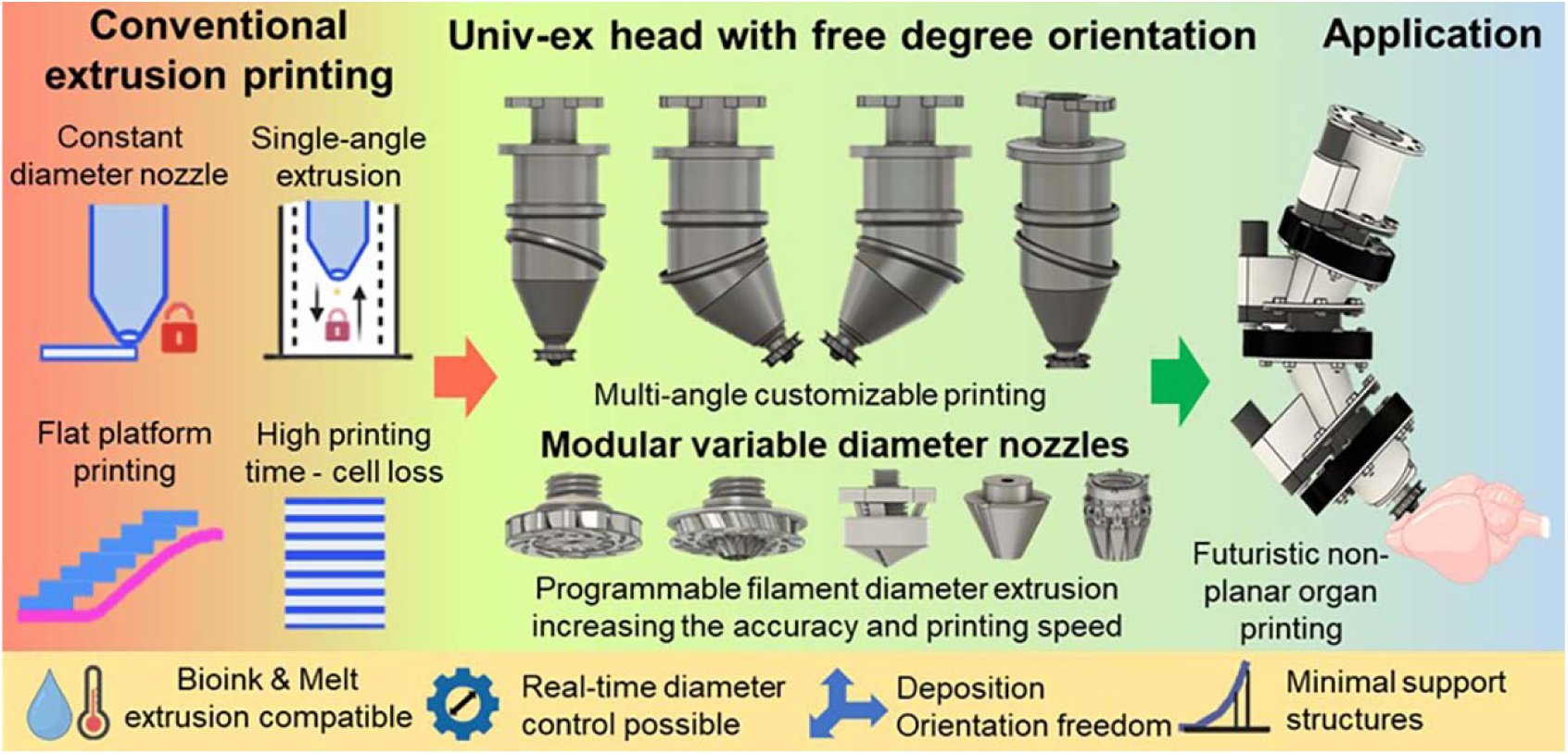

## 1. Introduction

Extrusion-based three-dimensional (3D) bioprinting has emerged as one of the most widely adopted bio-additive manufacturing strategies due to its material versatility, operational simplicity, and compatibility with a broad range of cell-laden and acellular bioinks [1–4]. It has enabled the fabrication of diverse tissue constructs for regenerative medicine, in vitro disease modeling, and biofabrication research.[5–7] Despite these advantages, extrusion-based systems remain fundamentally constrained by two long-standing hardware assumptions: fixed nozzle geometries and unidirectional, vertically aligned deposition. These constraints limit geometric adaptability and hinder the fabrication of conformal, non-planar, and physiologically relevant architectures.[8–11]

Fixed nozzle diameters impose a single resolution and flow regime throughout the printing process, making it challenging to fabricate constructs with spatially varying feature sizes such as branched vasculature, ducts, and graded tissue interfaces. Also, fixed deposition orientation confines extrusion primarily to planar, layer-by-layer strategies.[12–14] When applied to curved or organ-like geometries, this approach often leads to stair-stepping artifacts, void formation, reduced surface fidelity, and inefficient reliance on sacrificial support structures. Collectively, these limitations increase printing time, restrict design freedom, and may adversely affect biofabrication efficiency and construct fidelity.[15, 16]

To address these challenges, several strategies have been explored, including multi-nozzle systems, G-code modifications or T-codes (for tool changing), dynamic mixing approaches, filament diameter-adjustable 3D printing and adaptive extrusion mechanisms.[12–15, 17, 18] While these methods offer improvements in specific contexts, they often introduce additional complexity, require specialized nozzle fabrication, or lack unified adaptability in both deposition orientation and effective nozzle geometry. More recently, actuator-driven adaptive nozzle systems have demonstrated real-time modulation of filament dimensions through active diameter control[15]. Although promising, such systems typically rely on embedded actuators, compliant materials, or complex control architectures that can complicate miniaturization, sterilization, and translation to compact, manufacturable extrusion toolheads.[8, 15, 19] On the other hand, conformal and non-planar printing has been achieved using multi-axis robotic arms or complex gantry-based motion systems or modified G-codes.[15, 17–19] While effective, these approaches primarily shift complexity to the motion platform rather than addressing adaptability at the toolhead level. As a result, system-level degrees of freedom increase substantially, often at the expense of modularity, scalability, and broader adoption across existing extrusion-based bioprinting platforms.[4, 9]

In this context, there remains a critical need for hardware-level extrusion solutions that enable both orientation freedom and nozzle adaptability while preserving mechanical simplicity, manufacturability, and modular integration. Addressing these needs at the toolhead level - rather than through ink-specific strategies or system-wide motion complexity- offers a scalable pathway toward conformal bio-additive manufacturing. Here, we introduce a universal extrusion head (Univ-Ex head) that enables free-degree-of-orientation extrusion directly at the toolhead level through a mechanically simplified, swivel-based architecture. Unlike actuator-heavy or robot-assisted systems, the Univ-Ex head achieves orientation freedom through geometric design rather than active control in its present implementation, minimizing mechanical complexity while enabling deposition on angled and non-planar surfaces. In addition, we establish a modular nozzle ecosystem based on a standardized interface, providing a scalable foundation for adaptable and, in future implementations, diameter-tunable extrusion without reliance on soft-material deformation or complex actuator-driven mechanisms.

Importantly, this study is positioned as a hardware and manufacturing platform innovation, rather than a bioink-centric or biological validation study. To maintain rigorous and reproducible validation, the platform is intentionally evaluated using fixed-diameter nozzle variants, including flat and conical geometries, which provide stable numerical and experimental boundary conditions. Using a combination of parametric CAD design, stereolithography-fabricated prototypes, and fluid-dynamics simulations performed in FLOW-3D, we demonstrate stable filament formation across multiple nozzle diameters and establish the feasibility of angled and non-planar deposition. Furthermore, we introduce a gear-integrated Univ-Ex head architecture as a forward-looking design extension intended to enable synchronized orientation control and future G-code-driven actuation. Critically, experimentally validated components, numerically evaluated behaviors, and conceptual design upgrades are explicitly distinguished throughout the manuscript. This disciplined separation ensures transparency and aligns with the rigorous standards expected for platform-level innovations in advanced manufacturing. By localizing adaptability at the extrusion toolhead level, the Univ-Ex platform provides a manufacturable and scalable alternative to existing adaptive extrusion approaches. This work establishes a foundation for future integration of real-time control, metal-fabricated hardware, and closed-loop bio-additive manufacturing systems, with relevance across additive manufacturing, biofabrication, bioprinting, tissue engineering and regenerative medicine communities.

## 2. Materials and Methods

### 2.1 System Overview and Terminology

The extrusion platform comprises a universal extrusion head (Univ-Ex head) integrated with a modular nozzle ecosystem. The Univ-Ex head is implemented as a swivel-based extrusion head that enables free-degree orientation of the nozzle axis relative to the build plane through a passive mechanical joint. This architecture provides orientation freedom without the use of external actuators in the present prototype, thereby maintaining mechanical simplicity while enabling non-vertical deposition. In subsequent design iterations, gear elements are integrated into the Univ-Ex head to enable mechanically coupled, direction-specific rotation and synchronized orientation control, as conceptually demonstrated in later sections of this work.

The modular nozzle ecosystem is based on a standardized threaded interface that enables the rapid interchange of nozzle modules without modifying the main extrusion body. Two nozzle geometries were experimentally evaluated in this study: Nozzle 2, a flat-tip nozzle, and Nozzle 3, a conical nozzle. Each nozzle type was fabricated with fixed outlet diameters of 1.0, 1.5, and 2.0 mm to provide stable boundary conditions for numerical simulations and proof-of-concept extrusion experiments. However, a total of 5 nozzles types have been designed and integrated CAD design images are presented in the Supporting information. A gear-integrated Univ-Ex head variant was also developed as a forward-looking design intended to enable synchronized rotation and programmable orientation control. In the present study, this design was fabricated as a single-piece SLA prototype and evaluated for assembly feasibility and extrusion capability, without demonstrating synchronized multi-axis actuation during a single print.

### 2.2 Computer-Aided Design (CAD) of the Univ-Ex Head and Nozzle Modules

All components of the Univ-Ex head and modular nozzle system were designed using Autodesk Fusion 360. Parametric modeling was employed to iteratively refine key geometric parameters, including internal flow-channel geometry, nozzle outlet diameter, taper angle, bore length, and thread interfaces. Internal flow-contact regions were designed with smooth transitions to minimize dead volume and reduce abrupt shear gradients that could destabilize shear-thinning bioinks. The Univ-Ex head architecture was designed to support different angles relative to the build plane through a swivel-like joint geometry. Assembly interference checks and motion simulations were performed to verify fit, alignment, and rotational clearance across configurations. Final designs were exported as high-resolution, watertight STL files to preserve curved internal geometries relevant to flow modeling and fabrication. To enable integration with standard extrusion-based 3D bioprinters, a top connector compatible with syringe-based extrusion mounts was incorporated into the Univ-Ex head, allowing direct replacement of conventional syringe nozzles. This connector also provides an inlet for compressed-air or pressure-driven extrusion, which can be coupled to the printer’s existing pneumatic or pressure control system, and was evaluated for leak-free assembly and mechanical compatibility.

### 2.3 Prototype Fabrication by Stereolithography

Prototype components were fabricated using a Formlabs Form 3+/3B stereolithography printer with Clear Resin. This material was selected to enable visual inspection of internal channels, joint interfaces, and assembly tolerances during early-stage development. STL files were processed in PreForm software, and printing orientations were optimized to minimize supports within internal flow paths and threaded regions. A nominal layer thickness of 50 μm was used to balance geometric fidelity and fabrication time. Multiple Univ-Ex head prototypes, including fixed vertical and angled configurations, were assembled with flat and conical nozzle modules of 1.0, 1.5-, and 2.0-mm outlet diameters.

### 2.4 Post-Processing and Assembly

Printed components were washed in 99% isopropyl alcohol, air-dried, and post-cured according to manufacturer recommendations. Supports were carefully removed to avoid damage to nozzle outlets and sealing interfaces. Internal channels were flushed with solvent and compressed air to remove residual debris. Dimensional fit and rotational freedom were verified prior to assembly. Nozzle modules were threaded into the extrusion body, and the Univ-Ex head was assembled using the designed press-fit and threaded interfaces. Manual inspection confirmed smooth rotational motion and alignment across configurations. Custom-designed and 3D-printed connectors, dimensioned to be compatible with commonly used syringe-based extrusion mounts, were then used to integrate the assembled components into the printer; assemblies were evaluated for mechanical integrity and air-tightness under pressure-driven extrusion conditions.

### 2.5 Design Refinement and Downsizing (V1–V5)

V1–V5 denote successive design versions of the Univ-Ex extrusion head, where V represents version and each iteration reflects progressive geometric refinement and downsizing of the same core architecture. Initial Univ-Ex prototypes (V1) were intentionally oversized to accommodate robust implementation of the internal swivel mechanism and to simplify early assembly and visualization. Subsequent versions (V2–V5) focused on systematic downsizing and refinement to improve manufacturability, assembly reliability, and tolerance robustness under stereolithography (SLA) fabrication constraints. Joint clearances, snap-fit features, and friction-lock interfaces were iteratively adjusted to balance secure assembly with smooth rotational motion. Across iterations, the overall envelope of the extrusion head was reduced while preserving nozzle interchangeability and internal flow continuity. These refinements culminated in compact prototypes suitable for proof-of-concept extrusion testing and future translation to metal fabrication routes. In addition, a miniaturized Univ-Ex head prototype with an overall diameter of approximately 1 cm was fabricated as a proof of concept, demonstrating the feasibility of highly compact extrusion heads deployable for specific applications without external interchangeable nozzle modules.

### 2.6 Numerical Simulations

All numerical simulations were performed using FLOW-3D 2024R2 (Flow Science, Inc.) software. The incompressible Navier–Stokes equations were solved on a fixed Cartesian grid, with solid boundaries represented using the Fractional Area–Volume Obstacle Representation (FAVOR™) method. Critically, using a stationary Cartesian mesh avoids the need for remeshing as objects move and accurately captures complex nozzle features without the use of a body-fitted grid. Free-surface behavior was captured using the TruVOF approach.

A previously reported shear-thinning gelatin microgel ink, with rheological parameters selected to represent typical extrusion bioink behavior, was used for the simulation. Herschel–Bulkley constitutive model was used to describe the ink’s rheological behavior: *τ* = *τ*_0_ +*K γ̇ ^n^* with yield stress (*τ*_0)_) = 390.9 Pa, consistency index (*K*) = 128.3 Pa s *^n^*, and flow index (*n*) = 0.374, and density of 1070 kg m^-3^ [20].

Geometries were pre-processed to ensure watertight surfaces for numerical reconstruction. A single non-conforming mesh block was employed with an isotropic cell size of 0.425 mm. The top boundary condition was a pressure inlet, with an artificial solid mask restricting inflow to the nozzle orifice, similar to previously reported work [21, 22]. The bottom boundary was set as a moving wall with speeds of 0.1 cm/s□¹ for the 2.0 and 1.5 mm cases and 0.05 cm/s□¹ for the 1.0 mm case. Inlet pressures and substrate velocities were empirically tuned for each nozzle diameter to achieve stable filament formation. All other faces used symmetry or continuative conditions and were placed sufficiently far from the flow to avoid interference. No-slip was enforced at all fluid–solid interfaces. Filament widths were extracted from simulation outputs using ImageJ (Fiji), and statistical analysis was performed in OriginPro 2025b using two-way ANOVA followed by Tukey’s post hoc test; differences were considered not significant for p > 0.05.

### 2.7 Proof-of-Concept Extrusion Experiments

Proof-of-concept extrusion experiments were performed using fixed-diameter nozzle modules assembled with the Univ-Ex head. These experiments were designed to validate filament formation, nozzle compatibility, and diameter-dependent extrusion behavior. Simultaneous real-time diameter modulation and dynamic orientation changes were not implemented during a single experimental print. However, angled Univ-Ex head configurations were fabricated and assembled with different fixed-diameter nozzles to verify mechanical compatibility and extrusion feasibility under non-vertical orientations. A model hydrogel ink was prepared by dissolving 5 wt% sodium alginate in distilled water under heating and magnetic stirring until a homogeneous solution was obtained. Extrusion was performed using a RegenHU 3D Discovery bioprinter operated in pneumatic mode, in which the standard syringe head was replaced by the 3D-printed Univ-Ex head using custom-designed connectors. Flat (Nozzle 2) and conical (Nozzle 3) nozzle modules with outlet diameters of 1.0, 1.5, and 2.0 mm were evaluated. Larger-format Univ-Ex head prototypes were used for ease of handling during testing. Extruded filaments were imaged, and filament widths were quantified using ImageJ and analyzed in OriginPro 2025b.

**Scheme 1.**
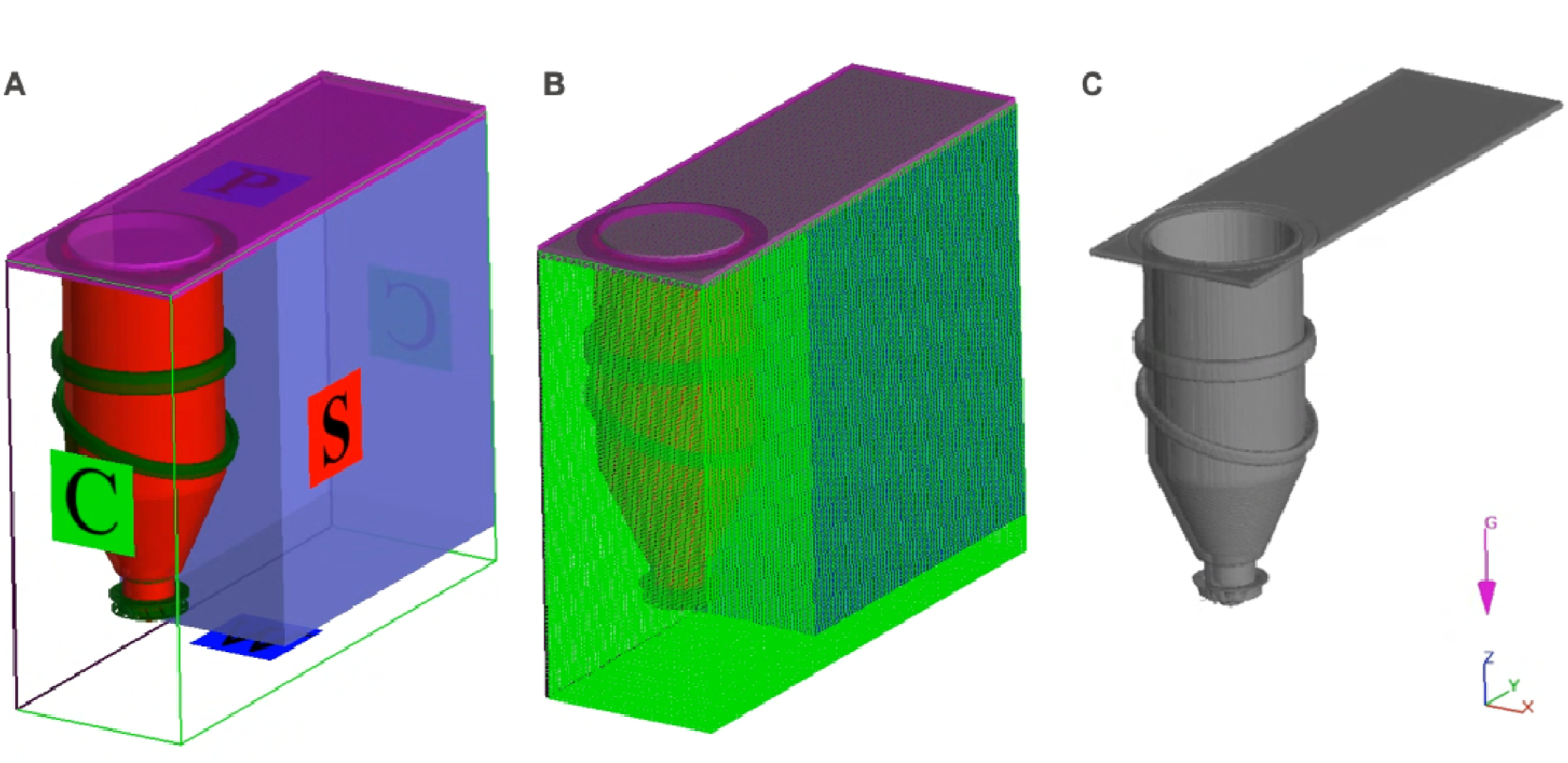
(A) Simulation setup in Flow3D. A solid wall on the top with a hole corresponding to the nozzle inlet. The bottom wall is moving and represents the build plate. The blue portion is a domain remover to decrease the number of cells. (B) Structured rectangular mesh generated by Flow3D. (C) Solid geometry generated by FAVOR™.

**Scheme 2.**
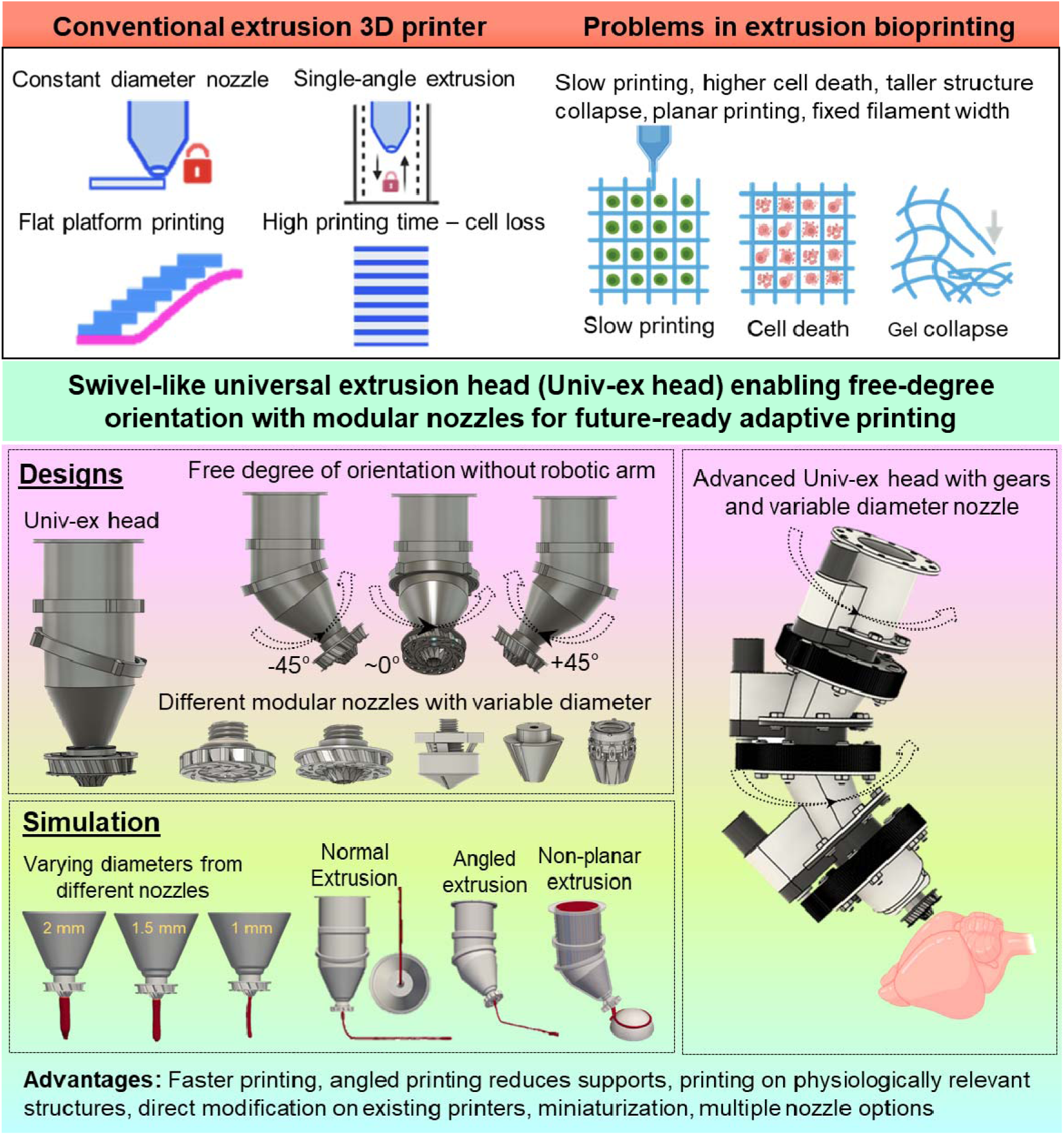
Summary of the concept: conventional planar extrusion (limitations) versus the Univ-Ex head enabling conformal and multiplanar deposition. Overview of the Univ-Ex extrusion platform, its working principle, and application scope. The scheme illustrates the concept of a free-degree-orientation universal extrusion (Univ-Ex) head with modular interchangeable nozzles, highlighting fixed-diameter extrusion in the present study and future-ready variable/gear-integrated concepts. Orientation-controlled extrusion enables conformal and non-planar deposition by coupling material extrusion with toolhead rotation and translational motion. The platform is applicable to extrusion-based bioprinting and bio-additive manufacturing, enabling improved filament placement on curved or complex geometries, reduced reliance on support structures, and compatibility with diverse printable materials for tissue engineering and advanced manufacturing applications.

## 3. Results and Discussion

### 3.1 Design rationale and positioning: overcoming fixed-diameter and fixed-orientation constraints

Extrusion-based bioprinting toolheads are fundamentally constrained by two hardware assumptions: that nozzle diameter remains constant throughout a print and that deposition occurs along a fixed, vertically aligned axis. These assumptions simplify motion control but significantly restrict geometric adaptability, particularly when printing heterogeneous or organ-inspired architectures. In practice, such constraints manifest as stair-stepping artifacts, void formation over curved surfaces, and inefficient reliance on sacrificial supports when approximating non-planar geometries. These limitations are well recognized in extrusion bioprinting and remain largely unresolved at the toolhead level. The Univ-Ex head was explicitly conceived to address these constraints through a toolhead-centric hardware strategy, rather than through increased gantry complexity or bioink-dependent solutions.[15, 17] As illustrated schematically in Scheme 2, conventional extrusion relies on planar layer stacking with a fixed nozzle orientation, whereas the proposed Univ-Ex head architecture enables multiplanar and conformal deposition by decoupling orientation freedom from the motion platform itself. Scheme 2 further contrasts experimentally validated core features with forward-looking design extensions, maintaining a clear separation between proof-of-concept implementation and future actuation capabilities.

Importantly, Scheme 2 distinguishes experimentally validated elements—a passive swivel-based orientation head with fixed-diameter nozzle modules—from forward-looking extensions, including active variable-diameter modulation and synchronized actuation, thereby maintaining strict claims discipline. This design philosophy differentiates the present work from prior approaches that either rely on multi-axis robotic arms to achieve conformality or introduce active actuators directly within the nozzle to alter filament dimensions.[4, 10, 15, 17] In contrast, the Univ-Ex head emphasizes mechanical simplification, modularity, and scalability, providing orientation freedom through geometry rather than complex control architectures.

### 3.2 Univ-Ex head architecture: mechanical simplification with orientation freedom

The Univ-Ex extrusion head was developed to address two persistent hardware constraints in extrusion-based bio-additive manufacturing: fixed deposition orientation and mechanically complex solutions for achieving angled or conformal printing. As illustrated in Figure 1a, the Univ-Ex head adopts a swivel-like joint architecture that enables free-degree orientation of the nozzle axis relative to the build plane while retaining a compact, toolhead-level design. Unlike robotic end-effector or multi-axis gantry approaches—where orientation control is achieved by adding degrees of freedom at the system level—the Univ-Ex concept embeds orientation adaptability directly within the extrusion head itself (**Supporting Videos 1-5**). This design philosophy minimizes mechanical overhead while remaining fully compatible with conventional extrusion platforms.

**Figure 1.**
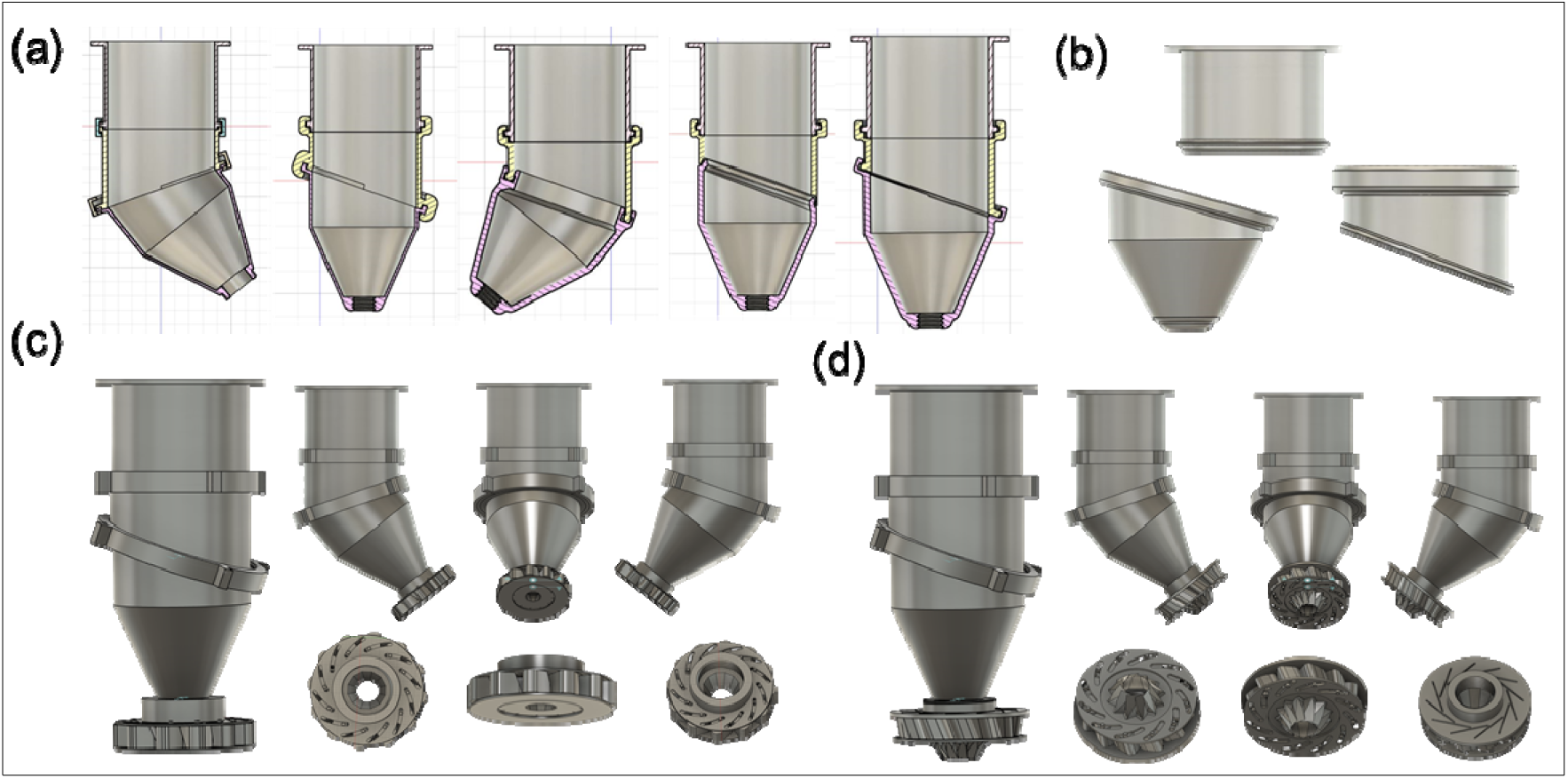
Design evolution and modular nozzle configurations of the Univ-Ex extrusion head. (a) Parametric CAD cross-section views illustrating the iterative design evolution of the swivel-based universal extrusion head (Univ-Ex head), highlighting variations in internal flow path geometry, joint inclination, and housing configuration across successive design iterations. (b) Exploded and isolated component views showing the modular architecture of the Univ-Ex head, including the interchangeable nozzle–housing interface and outlet components used for comparative evaluation. (c) Assembled Univ-Ex head equipped with the flat nozzle (Nozzle-3, flat outlet geometry), demonstrating passive swivel-enabled free-degree orientation using a fixed-diameter outlet. (d) Assembled Univ-Ex head equipped with the conical nozzle (Nozzle-3, conical outlet geometry), illustrating the same modular nozzle architecture and fixed-diameter configuration with an alternative outlet geometry.

The parametric CAD cross-section views in **Figure 1a** document the iterative design evolution of the Univ-Ex head, highlighting progressive refinements in internal flow-path continuity, joint inclination, and housing geometry. These iterations were guided by the need to preserve uninterrupted material flow during angular rotation while maintaining structural simplicity. Detailed engineering drawings, dimensional specifications, and sectional views provided in **Supporting Figures S1 and S2** further elucidate the internal channel geometry, swivel joint configuration, and standardized interface dimensions used throughout the study.

A key feature of the Univ-Ex architecture is its passive interlock-style swivel joint, which allows smooth angular rotation of the nozzle without the need for active actuation in the present implementation. This joint maintains a continuous internal flow path across a range of orientations, which is critical for extrusion stability and filament integrity during printing. From a mechanical standpoint, this approach avoids the cumulative positional errors, increased system footprint, and added control complexity typically associated with external multi-axis kinematics or robotic end-effector solutions.

The exploded and isolated component views in **Figure 1b** highlight the modular construction of the Univ-Ex head used for both experimental validation and numerical simulations. The standardized nozzle–housing interface enables rapid interchange of nozzle modules without altering the core extrusion head architecture. This modularity is central to the comparative evaluation of different outlet geometries and diameters while preserving identical upstream flow and boundary conditions.

**Figures 1c and 1d** show the assembled Univ-Ex head equipped with two validated nozzle configurations: a flat nozzle (Nozzle 2) and a conical nozzle (Nozzle 3), each operating with fixed outlet diameters. These assemblies demonstrate passive swivel-enabled free-degree orientation during angled printing while maintaining a mechanically simple, fixed-diameter extrusion strategy. Importantly, the same head architecture supports multiple nozzle geometries without modification, underscoring the flexibility of the modular design.

The validated modular nozzle designs and fixed outlet diameters used in simulation and experimental studies are summarized in **Supporting Figure S2**. By decoupling nozzle geometry and diameter selection from the core orientation mechanism, the Univ-Ex platform enables systematic investigation of diameter- and geometry-dependent extrusion behavior while maintaining consistent mechanical and kinematic conditions. Overall, the Univ-Ex head demonstrates that orientation freedom can be achieved through mechanical simplification rather than added system-level complexity. By preserving a low part count, compact form factor, and passive operation in its current form, the platform provides a robust and translation-ready foundation for future upgrades, including active actuation, synchronized control, and direct implementation in CNC-machined or metal additively manufactured toolheads for advanced bio-additive manufacturing applications.

### 3.3 Modular nozzle ecosystem: controlled validation using flat and conical geometries

Although different types (5 types – **Supporting Figure S3 and Supporting videos 1-5**) of nozzle concepts were explored during the design phase, the present study deliberately restricts experimental and numerical validation to two representative nozzle geometries: a flat nozzle (Nozzle 2) and a conical nozzle (Nozzle 3). This controlled selection was motivated by the need to limit design complexity, ensure reproducibility, and maintain stable and well-defined boundary conditions for both numerical simulations and experimental extrusion studies. The validated nozzle geometries and fixed outlet diameters are indicated in Figure 2.

**Figure 2.**
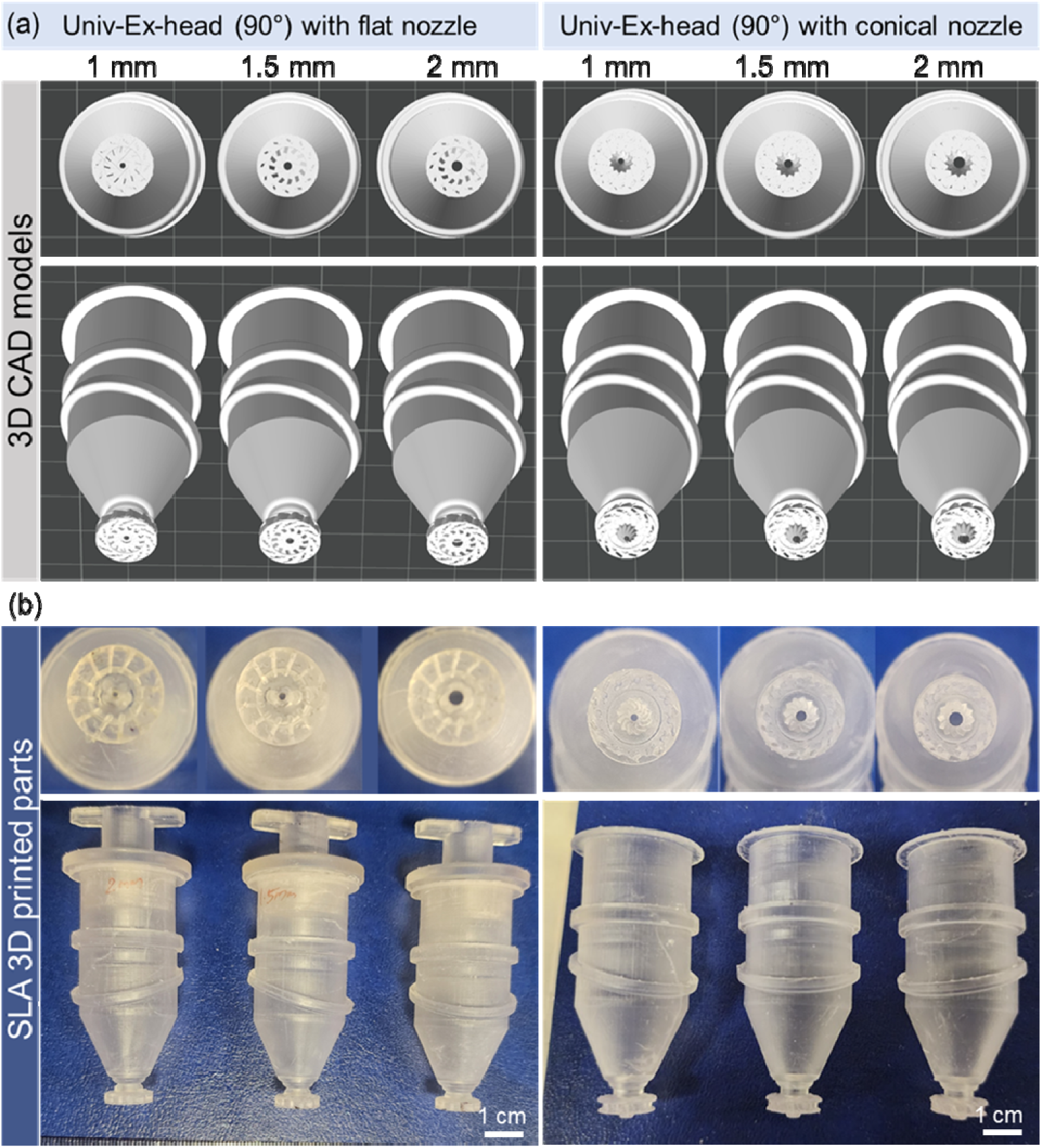
Assembly of the Univ-Ex head with modular nozzles and SLA-printed prototypes. (a) Sectional CAD views of the Univ-Ex head at 90° orientation, showing bottom and side views of interchangeable flat (Nozzle 2) and conical (Nozzle 3) nozzle geometries with fixed outlet diameters (1.0, 1.5, and 2.0 mm). The corresponding CAD assemblies illustrate the Univ-Ex head connected to each nozzle configuration. (b) Photographs of SLA-3D-printed prototypes (Clear Resin) showing fabricated components, assembled Univ-Ex head–nozzle units, and multiple nozzle diameters, demonstrating mechanical integrity, dimensional fidelity, and proof-of-concept assembly feasibility.

As shown in Figure 2a, both nozzle types were evaluated at identical orientations (90°) and fabricated with fixed outlet diameters of 1.0, 1.5, and 2.0 mm, enabling direct, geometry-dependent comparisons. The flat nozzle serves as a baseline outlet geometry, providing a uniform exit profile and predictable filament footprint under planar and angled deposition conditions. In contrast, the conical nozzle incorporates a tapered outlet region, which modifies local shear distribution and flow convergence near the nozzle exit. This geometric variation enables systematic assessment of how outlet shape influences filament formation, continuity, and diameter stability under otherwise identical extrusion conditions.

Crucially, both nozzle geometries were implemented as modular, interchangeable components using a standardized threaded interface, as demonstrated by the assembled and standalone parts in Figure 2b. This modular architecture decouples nozzle geometry from the extrusion head body, representing a mechanical advantage over monolithic nozzle designs reported in earlier studies, where changing outlet geometry often necessitates redesign or re-fabrication of the entire extrusion assembly.[10, 23, 24] In the Univ-Ex platform, nozzle interchangeability enables rapid reconfiguration without altering the core head architecture, supporting systematic validation in the present study and facilitating future design evolution toward additional geometries or actively tunable nozzle concepts.

### 3.4 Prototype realization and mechanical fidelity of SLA-fabricated components

The manufacturability and assembly fidelity of the Univ-Ex head were evaluated using SLA-fabricated prototypes in Clear Resin. As shown in Figure 2, sectional CAD views, assembled renderings, and photographs of printed components collectively demonstrate accurate reproduction of internal channels, swivel joints, and threaded interfaces. The transparency of the printed parts enabled direct inspection of flow pathways and verification of internal continuity after assembly. From a mechanical validation perspective, the SLA prototypes confirm several critical aspects of the design. First, the swivel joint maintains smooth rotational motion while preserving alignment between the nozzle outlet and the internal flow path. Second, the standardized nozzle interface supports repeated installation and removal without degradation of fit or alignment. Third, downsizing iterations (V1–V5) successfully reduced the overall envelope of the extrusion head while maintaining tolerance robustness under SLA fabrication constraints. It is important to emphasize that the SLA-fabricated prototypes presented in this study represent development-stage hardware intended to validate geometry, assembly logic, and mechanical feasibility, rather than long-term wet-use performance. The manuscript makes no claims regarding durability, sterilizability, or biocompatibility of the Clear Resin components. The Univ-Ex head architecture is intentionally designed to be material- and process-agnostic, enabling straightforward future translation to CNC-machined or metal additively manufactured (e.g., aluminum or steel) toolheads, where tighter tolerances, improved surface finish, and enhanced operational robustness can be achieved.

### 3.5 Numerical–experimental validation of extrusion behavior across nozzle diameters

FLOW-3D simulations were employed as a central validation tool to quantify extrusion behavior across nozzle geometries and outlet diameters. The simulation framework, incorporating FAVOR™-based geometry reconstruction and TruVOF free-surface tracking, enables resolution of filament formation dynamics under controlled inlet pressure and substrate motion conditions. As shown in **Figure 3a**, the simulations predict stable filament formation across fixed outlet diameters of 1.0, 1.5, and 2.0 mm for both nozzle types (Nozzle 2 & 3). Analysis of the simulated velocity fields and free-surface evolution (**Figure 3a**) reveals clear diameter-dependent flow regimes, with larger outlets producing broader filaments and reduced sensitivity to local perturbations, while smaller outlets exhibit tighter confinement and increased shear localization near the nozzle exit.

**Figure 3.**
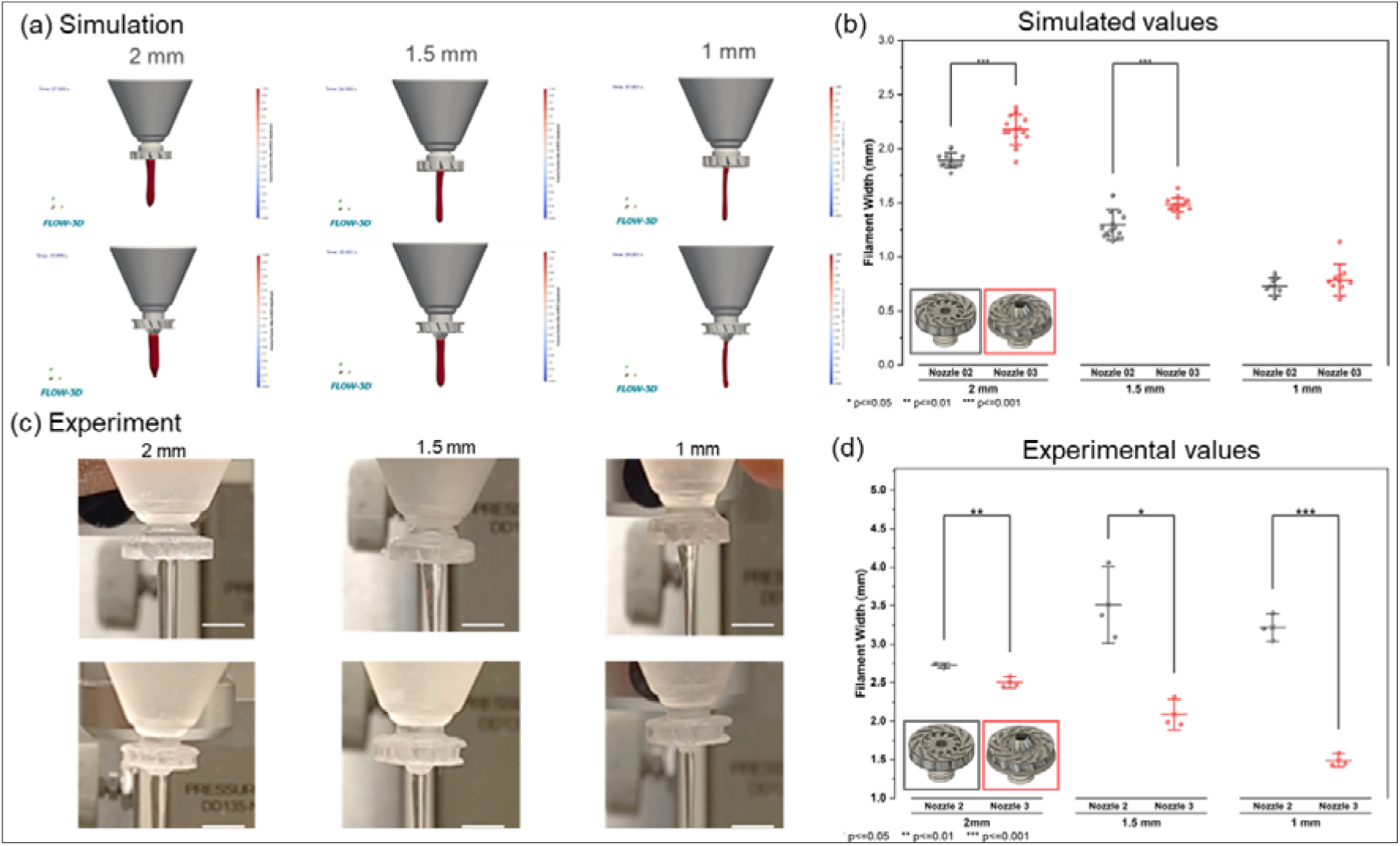
Simulation–experiment validation of filament extrusion using fixed-diameter nozzles. (a) FLOW-3D simulation snapshots showing filament formation and free-surface evolution during extrusion through Nozzle 2 (flat) and Nozzle 3 (conical) with fixed outlet diameters of 2.0, 1.5, and 1.0 mm. (b) Quantitative filament widths extracted from the simulation results using ImageJ, comparing Nozzle 2 and Nozzle 3 across different outlet diameters and demonstrating diameter-dependent extrusion behavior. (c) Corresponding experimental images of alginate-based gel extrusion obtained using stereolithography-printed Univ-Ex prototypes with fixed outlet diameters of 2.0, 1.5, and 1.0 mm for both nozzle geometries. (d) Experimentally measured filament widths quantified from optical images using ImageJ, highlighting reliable diameter-dependent trends.

Quantitative filament widths extracted from the numerical results (**Figure 3b**) show systematic, diameter-dependent trends for both flat (Nozzle 2) and conical (Nozzle 3) geometries. These results provide a comparative numerical baseline for evaluating outlet geometry and diameter effects under identical boundary conditions. Corresponding proof-of-concept extrusion experiments (**Figure 3c**) demonstrate filament formation that follows the same diameter-dependent trends predicted numerically. Experimentally measured filament widths (**Figure 3d**) decrease consistently with reducing outlet diameter for both nozzle geometries. This agreement supports two key conclusions. First, the CAD-defined nozzle geometries are faithfully reconstructed within the numerical framework, validating the simulation approach for comparative design screening. Second, the numerical model provides a rational basis for selecting inlet pressures and substrate velocities that sustain continuous filament formation, reducing reliance on exhaustive trial-and-error experimentation. Compared to earlier extrusion studies that rely primarily on empirical tuning, this combined simulation–experiment strategy strengthens the mechanical design narrative and enables predictive optimization prior to metal fabrication.[25, 26]

### 3.6 Diameter-dependent extrusion patterns under planar deposition - simulation

To establish a baseline understanding of extrusion behavior prior to orientation-controlled studies, FLOW-3D simulations were first used to examine planar filament deposition across different nozzle outlet diameters. As shown in Figure 4a, side-view simulations capture stable filament formation during extrusion for fixed outlet diameters of 2.0, 1.5, and 1.0 mm, with continuous strands maintained under constant inlet pressure and substrate translation. The corresponding bottom-view simulations (Figure 4b) provide direct visualization of filament footprint, width uniformity, and deposition trajectory on the build surface. Clear diameter-dependent trends are observed, with larger outlet diameters producing broader filaments and smaller outlets yielding narrower, more confined deposition paths. These results confirm that the numerical framework accurately captures the influence of nozzle diameter on filament morphology under planar printing conditions. Importantly, these planar simulations serve as a reference condition for subsequent orientation-controlled and non-planar deposition studies presented in the following section. By first validating diameter-dependent extrusion behavior under well-defined planar motion, the simulations establish a reliable foundation for extending the analysis to angled and conformal deposition scenarios using the same numerical framework.

**Figure 4.**
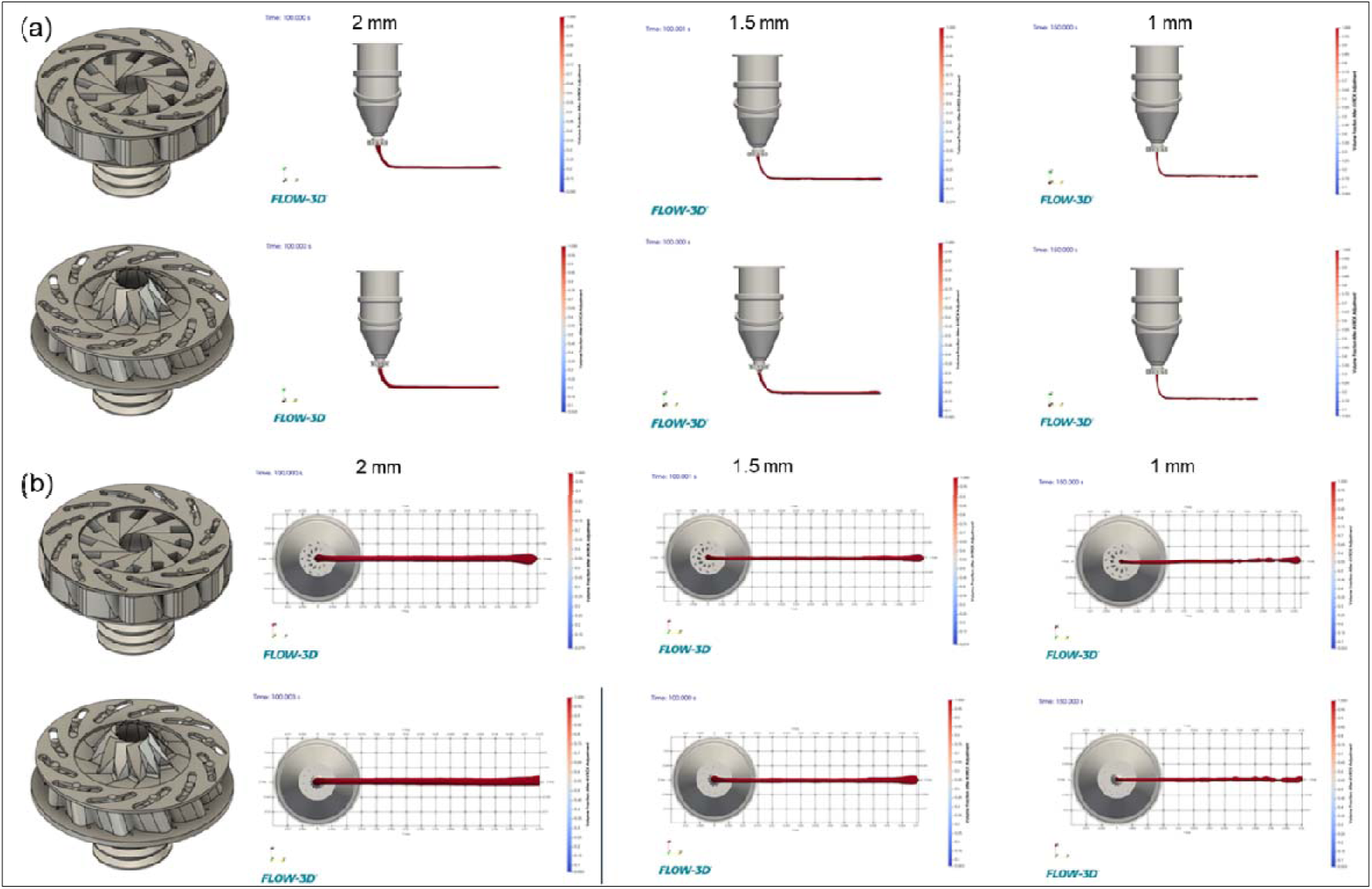
Numerical simulation of filament extrusion and deposition through different nozzle diameters using FLOW-3D. (a) Side-view snapshots of the extrusion process showing continuous filament formation and deposition during printing for nozzle outlet diameters of 2.0, 1.5, and 1.0 mm. The simulations capture the coupled effects of material extrusion and relative motion between the nozzle and the moving build surface, illustrating filament bending, stabilization, and laydown behavior along the printing direction. (b) Corresponding bottom-view snapshots of the same simulations, highlighting filament width evolution, continuity, and trajectory during deposition for each nozzle diameter. Together, the side and bottom views demonstrate the ability of the numerical model to resolve diameter-dependent extrusion behavior and dynamic filament deposition under printing-relevant conditions.

### 3.7 Orientation-controlled extrusion and feasibility of non-planar deposition

A defining capability of the Univ-Ex head is its mechanically enabled orientation freedom, which provides a direct pathway toward conformal deposition on non-planar surfaces. While dynamic orientation control during a single experimental print was not demonstrated in the present study, FLOW-3D simulations provide compelling evidence for the feasibility of orientation-controlled extrusion. As shown in Figure 5a, numerical simulations demonstrate stable filament formation during angled extrusion across fixed nozzle outlet diameters of 2.0, 1.5, and 1.0 mm (**Supporting videos 6-12**). The results indicate that inclined nozzle orientation does not inherently destabilize extrusion, provided that inlet pressure and translational motion are appropriately tuned. Diameter-dependent filament behavior observed under planar conditions is preserved under angled extrusion, confirming that orientation changes can be accommodated within the same flow regime.

**Figure 5.**
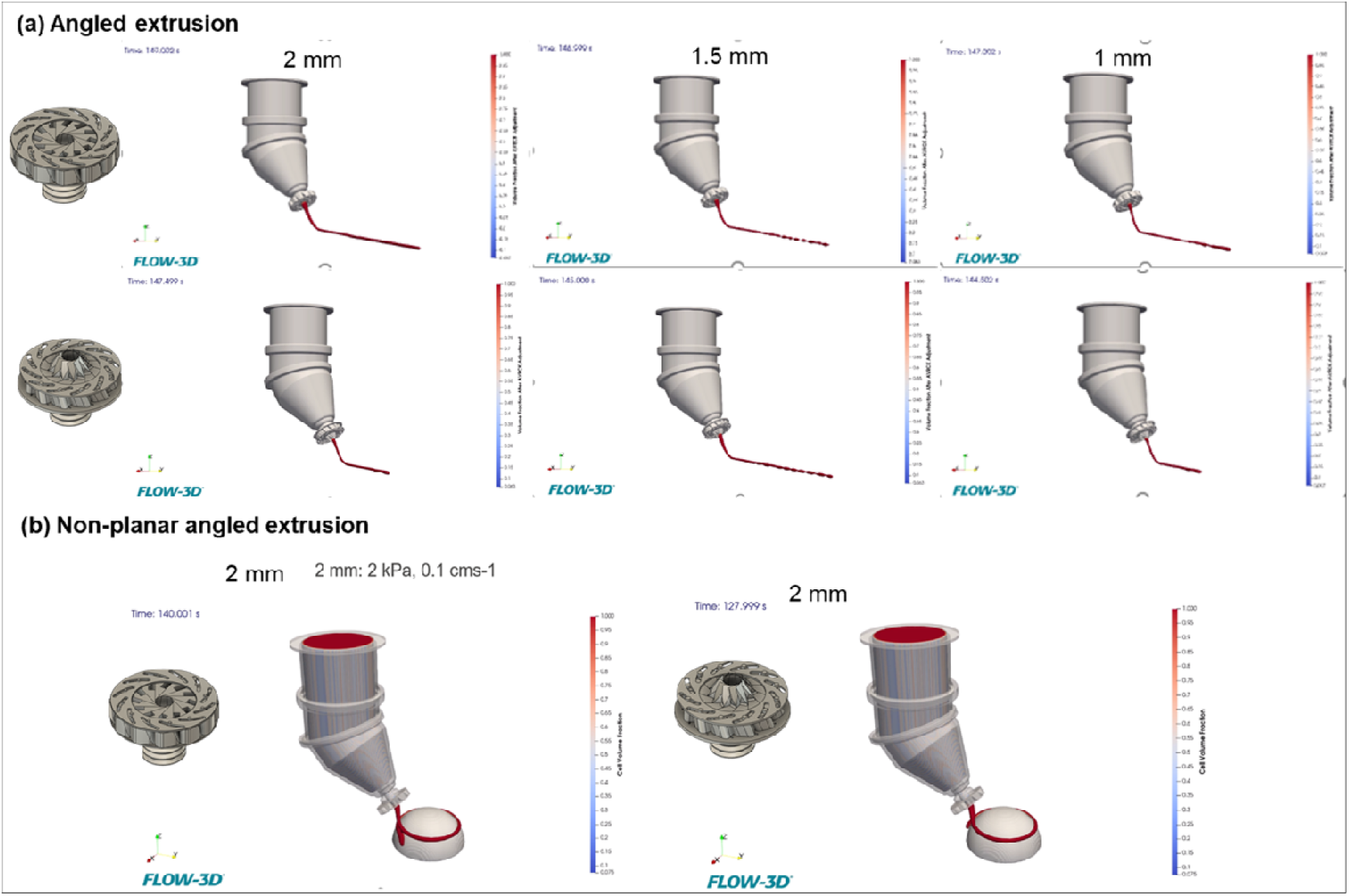
Diameter- and orientation-dependent extrusion behavior visualized through FLOW-3D simulations. (a) Side-view snapshots of filament extrusion from 2.0, 1.5, and 1.0 mm nozzle outlets during simultaneous material extrusion and translational motion, illustrating stable filament formation, curvature, and laydown behavior as a function of nozzle diameter. Color contours indicate the volume fraction of the extruded material, capturing continuous filament generation under printing-relevant conditions. (b) Three-dimensional simulation views demonstrating extrusion onto a non-planar substrate with the nozzle oriented at an inclined angle relative to the build platform, highlighting coupled extrusion and motion during angled printing. These results illustrate the capability of the numerical framework to resolve filament deposition during orientation-controlled printing on curved or non-planar surfaces.

Extending this analysis, Figure 5b illustrates non-planar angled extrusion, where continuous filaments are deposited onto a curved substrate under inclined nozzle orientation. These simulations show that conformal deposition can be achieved without fundamentally altering extrusion physics, but rather by enabling mechanical orientation freedom at the toolhead level. The modeled flow fields and free-surface evolution confirm that continuous strand formation can be maintained along non-planar deposition pathways. Experimentally, angled Univ-Ex head configurations were fabricated and assembled with different nozzle modules to verify mechanical compatibility under non-vertical orientations. Although these tests do not constitute dynamic orientation-controlled printing, they confirm that the extrusion head geometry, swivel interface, and modular nozzle connections remain mechanically viable when oriented away from the vertical axis.

Relative to prior approaches that rely on complex robotic motion platforms to achieve conformal printing, the Univ-Ex head offers a complementary, toolhead-centric strategy in which orientation adaptability is embedded directly within the extrusion head. Collectively, the numerical and experimental observations establish that: (i) the Univ-Ex head geometry is mechanically compatible with angled deposition, (ii) the flow physics supports stable extrusion under angled and non-planar trajectories, and (iii) orientation-controlled extrusion represents a justified pathway for future integration with toolpath planning, synchronized actuation, and closed-loop motion control.

### 3.8 Gear-integrated Univ-Ex head: mechanical pathway toward synchronized actuation

To explore future integration of synchronized orientation control within the Univ-Ex platform, a gear-integrated variant of the extrusion head was developed as a conceptual mechanical upgrade. The proposed architecture and component layout are shown in Figure 6a–d, where integrated gear coupling and a slew-bearing interface are designed to enable mechanically synchronized rotation between head segments, providing a pathway toward future G-code-controlled actuation. As a proof-of-feasibility step, the gear-integrated Univ-Ex head was fabricated as a single-piece SLA prototype and assembled with both flat (Nozzle 2) and conical (Nozzle 3) modular nozzle configurations across fixed outlet diameters (Figure 6e,f). Successful filament extrusion from this prototype confirms that the added geometric and mechanical complexity introduced by the gear architecture does not preclude continuous filament formation, supporting the mechanical viability of the concept. Importantly, synchronized multi-axis actuation during a single print was not implemented in the present work. Accordingly, the gear-integrated Univ-Ex head is treated strictly as a forward-looking design upgrade, rather than a fully validated functional system. Overall, the gear-integrated design establishes a mechanically plausible pathway for future developments incorporating programmed orientation control, synchronized rotation, and closed-loop actuation, building directly upon the passive swivel-based Univ-Ex architecture validated in this study.

**Figure 6.**
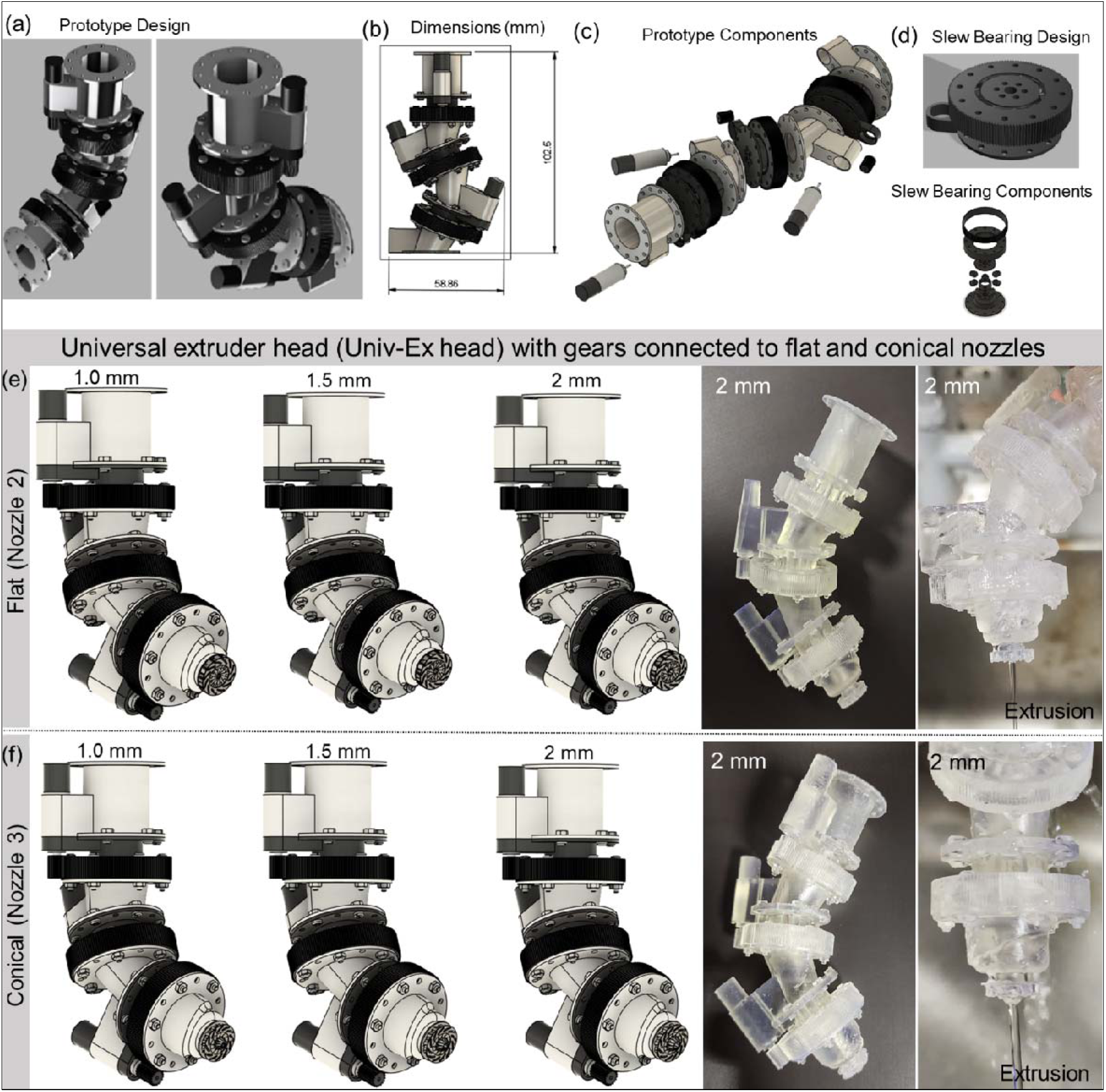
Gear-integrated Univ-Ex head (conceptual upgrade) CAD renderings and SLA 3D printed prototypes. (a) CAD rendering of the gear-integrated Univ-Ex head illustrating the conceptual design intended for synchronized multi-axis actuation. (b) CAD view showing the overall dimensions of the gear-integrated Univ-Ex head. (c) Exploded CAD view highlighting the internal gear features and constituent components. (d) Detailed CAD views of the slew bearing design and its components. (e) CAD views of the single-piece gear-integrated Univ-Ex head, along with the assembled prototype fitted with flat (Nozzle 2) interchangeable nozzle modules (1.0, 1.5, and 2.0 mm outlet diameters). (f) CAD and photographic views of the assembled prototype fitted with conical (Nozzle 3) nozzle modules, including photographs of the single-piece SLA-printed gear-integrated Univ-Ex head (2.0 mm nozzle) and a proof-of-concept extrusion demonstration.

### 3.9 Comparison with prior extrusion and orientation-control strategies

Several recent academic studies have explored adaptive extrusion strategies to overcome the fixed-resolution limitations of conventional material extrusion printing. Kang and Mueller introduced adaptive nozzle 3D printing (AN3DP), where nozzle diameter and cross-sectional shape are actively modulated in real time using a tendon-driven pin array and compliant membrane, enabling continuous multiscale and gradient fabrication.[15] Similarly, Qu et al. demonstrated filament diameter–adjustable printing by dynamically varying printing velocity and nozzle–substrate distance, enabling complex 1D–3D gradient structures within conventional direct ink writing systems.[17] Other reported studies have investigated rotating or non-circular nozzle apertures to influence filament geometry and deposition behavior, using nozzle rotation or aperture shape to tune bead width and morphology. Collectively, these approaches primarily achieve adaptive extrusion through active actuation, process-parameter modulation, or nozzle rotation, while generally maintaining fixed deposition orientation. In contrast, the Univ-Ex platform explored in this work introduces a toolhead-level orientation strategy, enabling angled and non-planar deposition independent of filament diameter modulation, and thereby occupying a complementary design space within extrusion-based additive manufacturing (**Supporting information Table S1**).

## 4. Limitations and Future Work

This study is intentionally design-forward and hardware-focused. Real-time variable-diameter extrusion and synchronized orientation control during a single print were not experimentally implemented; instead, fixed-diameter nozzles and passive orientation were used to maintain stable boundary conditions, with numerical simulations supporting feasibility of angled and non-planar deposition. The SLA-printed prototypes represent development-stage hardware for geometric and mechanical validation only, with no claims regarding long-term durability, sterilizability, or biocompatibility. Future work will focus on metal-fabricated toolheads, integration of active actuation for synchronized control, expansion of the modular nozzle library, and coupling toolhead-level orientation with advanced toolpath planning and closed-loop control.

## 5. Conclusions

This work introduces a universal extrusion head (Univ-Ex head) and modular nozzle platform designed to address two persistent hardware bottlenecks in extrusion bioprinting: fixed nozzle orientation and limited nozzle adaptability. The Univ-Ex head enables free-degree-of-orientation extrusion through a mechanically simplified swivel architecture, while a standardized modular interface allows rapid interchange of fixed-geometry nozzle modules. Validation combining FLOW-3D simulations and SLA-printed development prototypes demonstrates stable filament extrusion across fixed outlet diameters and supports the feasibility of angled and non-planar deposition trajectories. A gear-integrated Univ-Ex design is presented as a forward-looking mechanical upgrade and demonstrated as a single-piece SLA proof-of-concept to confirm extrusion feasibility without synchronized actuation. Collectively, these results establish a scalable, toolhead-centric hardware pathway toward conformal bio-additive manufacturing and motivate future translation to metal-fabricated systems with programmable orientation control and, where appropriate, real-time diameter modulation.

## Acknowledgements

We thank the NYU Abu Dhabi Core Technology (NYUAD CTP) Platform for providing access to instrumentation. We also acknowledge the assistance of undergraduate students Guglielmo Fonda, Julio Zuazola, Selina Shah and Abbad Shazly in supporting this work.

## AI Disclosure Statement

During the preparation of this work, the author(s) used ChatGPT-5.2 (OpenAI) to assist with grammar correction and language editing. After using this tool, the author(s) carefully reviewed and edited the content as needed and take full responsibility for the content of the published article.

## Contributions

G.J. and S.V. conceived the study. G.J. designed the research, prepared the inks, developed the design-to-fabrication workflow, fabricated the prototypes, acquired images and videos, and analyzed the data. R.C. performed the numerical simulations. G.J. wrote the manuscript with input from all authors. S.V. supervised the project.

## Funding

This research received no external grant from any funding agency in the public, commercial, or not-for-profit sectors.

## Declaration of interest statement

G.J. and S.V. have filed a provisional U.S. patent application related to the technology described in this manuscript. R.C. declares no competing intellectual property interests.

## Data availability

All relevant data supporting the findings of this study are available within the article and its Supporting Information files as well as Source Data. Additional datasets are available from the corresponding author upon reasonable request. Source data are provided in this paper.

## Notes

### Competing Interest Statement

SV and GJ have filed a provisional U.S. patent application related to the technology described in this manuscript. Author RC declares no competing intellectual property interests.

## References

[1] J. Gopinathan, I. Noh, Recent trends in bioinks for 3D printing, Biomater Res 22 (2018) 11.

[2] S.V. Murphy, A. Atala, 3D bioprinting of tissues and organs, Nat Biotechnol 32(8) (2014) 773–85.

[3] S. Vijayavenkataraman, W.C. Yan, W.F. Lu, C.H. Wang, J.Y.H. Fuh, 3D bioprinting of tissues and organs for regenerative medicine, Adv Drug Deliv Rev 132 (2018) 296–332.

[4] Y.S. Zhang, G. Haghiashtiani, T. Hübscher, D.J. Kelly, J.M. Lee, M. Lutolf, M.C. McAlpine, W.Y. Yeong, M. Zenobi-Wong, J. Malda, 3D extrusion bioprinting, Nature Reviews Methods Primers 1(1) (2021) 75.

[5] L. Moroni, J.A. Burdick, C. Highley, S.J. Lee, Y. Morimoto, S. Takeuchi, J.J. Yoo, Biofabrication strategies for 3D in vitro models and regenerative medicine, Nature Reviews Materials 3(5) (2018) 21–37.

[6] M. Ryma, T. Scheibel, Bridging the Gap: Unlocking the Potential of Biofabrication for Applications in In Vitro Testing, Langmuir 41(31) (2025) 20433–20442.

[7] M.A. Heinrich, W.J. Liu, A. Jimenez, J.Z. Yang, A. Akpek, X. Liu, Q.M. Pi, X. Mu, N. Hu, R.M. Schiffelers, J. Prakash, J.W. Xie, Y.S. Zhang, 3D Bioprinting: from Benches to Translational Applications, Small 15(23) (2019).

[8] I.T. Ozbolat, M. Hospodiuk, Current advances and future perspectives in extrusion-based bioprinting, Biomaterials 76 (2016) 321–343.

[9] I. Holland, Extrusion bioprinting: meeting the promise of human tissue biofabrication?, Prog Biomed Eng 7(2) (2025).

[10] L. Lombardi, A. Scalzone, C. Ausilio, P. Gentile, D. Tammaro, Optimizing nozzle design in extrusion-based 3D bioprinting to minimize mechanical stress and enhance cell viability, IJB 11(4) (2025).

[11] A.A. Armstrong, J. Norato, A.G. Alleyne, A.J.W. Johnson, Direct process feedback in extrusion-based 3D bioprinting, Biofabrication 12(1) (2020).

[12] T. Tezel, V. Kovan, Determination of optimum production parameters for 3D printers based on nozzle diameter, Rapid Prototyping J 28(1) (2022) 185–194.

[13] J. Wang, T.W. Chen, Y.A. Jin, Y. He, Variable bead width of material extrusion-based additive manufacturing, J Zhejiang Univ-Sc A 20(1) (2019) 73–82.

[14] T. Kuipers, E.L. Doubrovski, J. Wu, C.C.L. Wang, A Framework for Adaptive Width Control of Dense Contour-Parallel Toolpaths in Fused Deposition Modeling, Comput Aided Design 128 (2020).

[15] S.W. Kang, J. Mueller, Multiscale 3D printing via active nozzle size and shape control, Sci Adv 10(23) (2024).

[16] A. Rossi, T. Pescara, A.M. Gambelli, F. Gaggia, A. Asthana, Q. Perrier, G. Basta, M. Moretti, N. Senin, F. Rossi, G. Orlando, R. Calafiore, Biomaterials for extrusion-based bioprinting and biomedical applications, Front Bioeng Biotech 12 (2024).

[17] H.W. Qu, C.J. Gao, K.Z. Liu, H.Y. Fu, Z.Y. Liu, P.H.J. Kouwer, Z.Y. Han, C.S. Ruan, Gradient matters via filament diameter-adjustable 3D printing, Nat Commun 15(1) (2024).

[18] S. Propst, J. Mueller, Time Code for multifunctional 3D printhead controls, Nat Commun 16(1) (2025).

[19] N.C. Brown, D.C. Ames, J. Mueller, Multimaterial extrusion 3D printing printheads, Nature Reviews Materials 10(11) (2025) 807–825.

[20] K. Song, D. Zhang, J. Yin, Y. Huang, Computational study of extrusion bioprinting with jammed gelatin microgel-based composite ink, Additive Manufacturing 41 (2021) 101963.

[21] K. El Abbaoui, I. Al Korachi, M. El Jai, B. Šeta, M.T. Mollah, 3D concrete printing using computational fluid dynamics: Modeling of material extrusion with slip boundaries, Journal of Manufacturing Processes 118 (2024) 448–459.

[22] M.T. Mollah, R. Comminal, W.R. Leal da Silva, B. Šeta, J. Spangenberg, Computational fluid dynamics modelling and experimental analysis of reinforcement bar integration in 3D concrete printing, Cement and Concrete Research 173 (2023) 107263.

[23] P.J. Mccauley, A.V. Bayles, Nozzle Innovations That Improve Capacity and Capabilities of Multimaterial Additive Manufacturing, Acs Eng Au 4(4) (2024) 368–380.

[24] Y.Y. Xu, C.J. Wang, Y. Yang, H. Liu, Z. Xiong, T. Zhang, W. Sun, A Multifunctional 3D Bioprinting System for Construction of Complex Tissue Structure Scaffolds: Design and Application, Int J Bioprinting 8(4) (2022) 254–273.

[25] J.C.G. Blanco, A. Macías-García, J.M. Rodríguez-Rego, L. Mendoza-Cerezo, F.M. Sánchez-Margallo, A.C. Marcos-Romero, J.B. Pagador-Carrasco, Optimising Bioprinting Nozzles through Computational Modelling and Design of Experiments, Biomimetics-Basel 9(8) (2024).

[26] R. Chand, B.S. Muhire, S. Vijayavenkataraman, Computational Fluid Dynamics Assessment of the Effect of Bioprinting Parameters in Extrusion Bioprinting, Int J Bioprinting 8(2) (2022) 45–60.

