## Supplementary material for "A Universal Free-Degree Orientation Extrusion Head Enables Conformal and Non-Planar Bio-Additive Manufacturing toward Adaptive and Future-Ready Bioprinting": Supporting information - Nozzle Univ-Ex head manuscript.docx

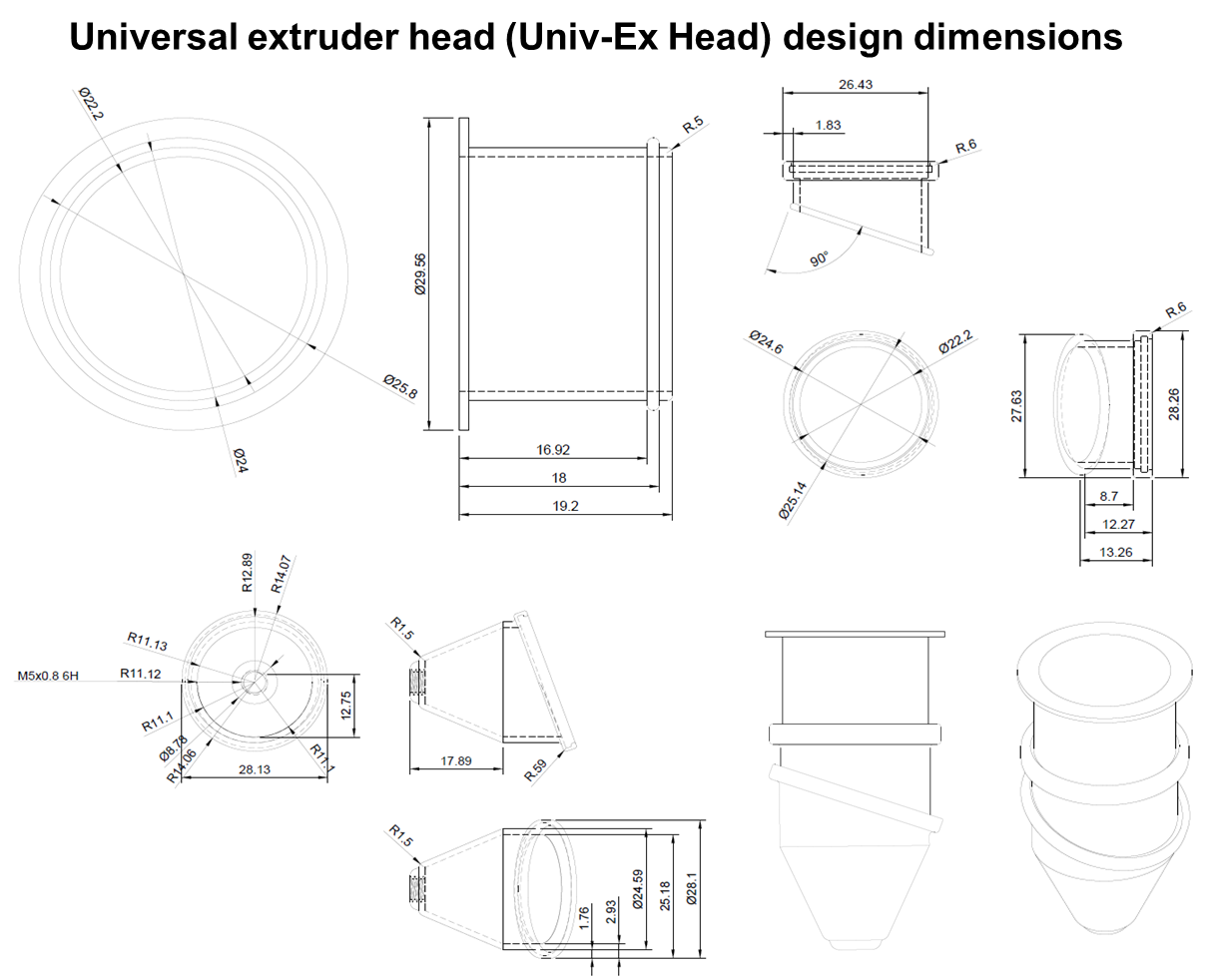

**Figure S1. Dimensional specifications of the universal extrusion head (Univ-Ex head).**
Annotated CAD views showing the key geometric dimensions of the Univ-Ex head, including overall height, housing diameter, swivel joint length, nozzle interface dimensions, and outlet positioning. All dimensions correspond to the proof-of-concept polymeric prototypes used for numerical simulations and experimental validation in this study.

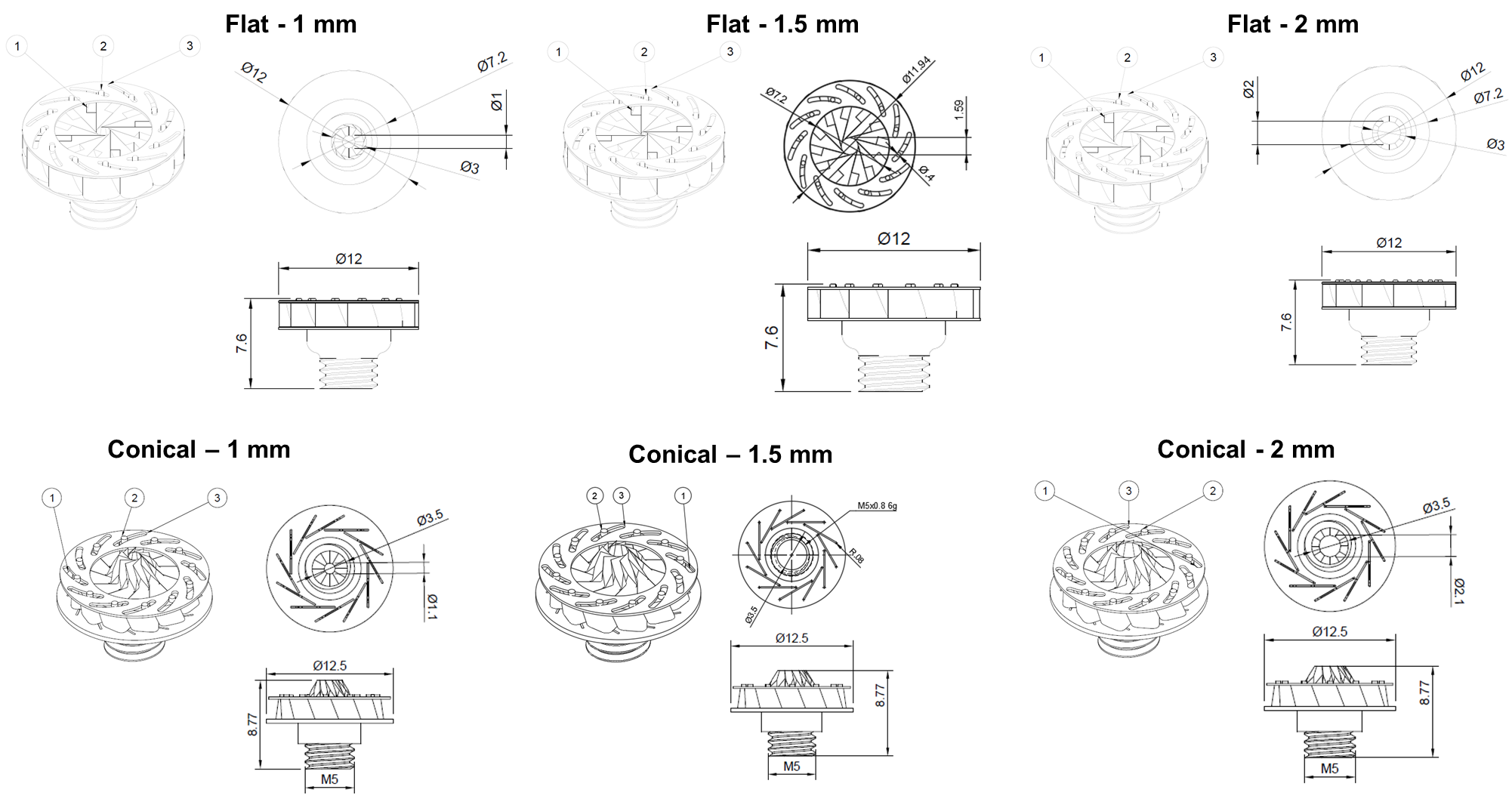

**Figure S2.** Computer-aided design (CAD) drawings of the modular nozzle architectures used in this study. The figure shows flat-tip and conical-tip nozzle designs with nominal outlet diameters of 1.0, 1.5, and 2.0 mm, including top, side, and sectional views with key dimensional annotations. These interchangeable nozzle geometries were designed in a parametric manner to enable systematic evaluation of diameter-dependent extrusion behavior and to ensure compatibility with the universal swivel-based extrusion head used for experimental validation and numerical simulations.

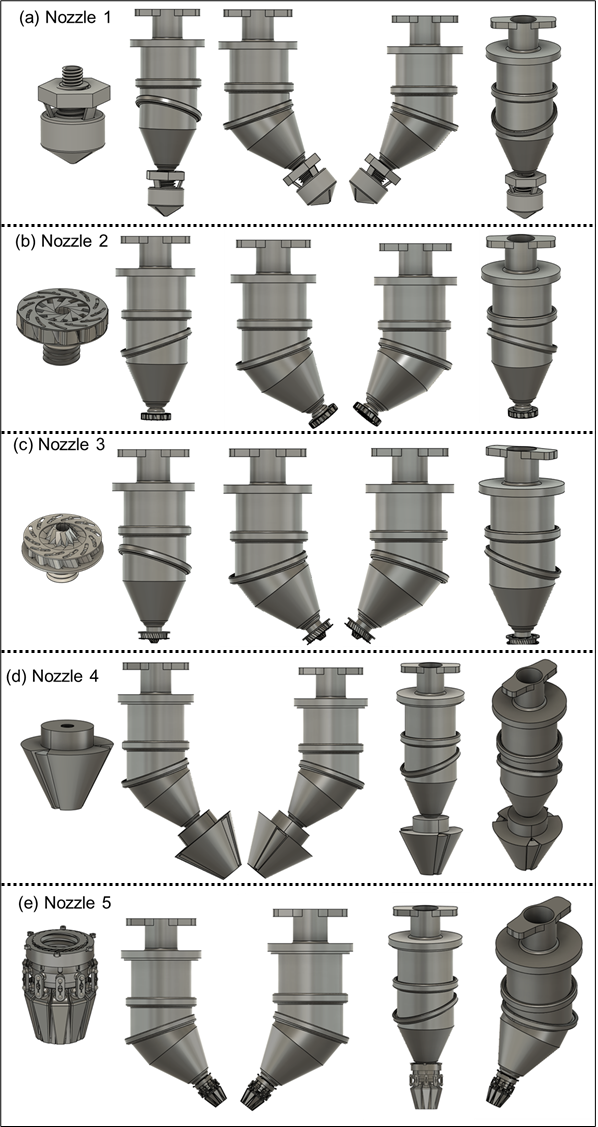

**Figure S3.** CAD representations of the modular nozzle assembly with universal swivel-based extrusion head designed in this study. The figure illustrates the overall geometry, external features, and interchangeable nozzle interface designed to enable free-degree orientation and compatibility with multiple nozzles for extrusion-based printing.

Table S1: Comparison of different variable filament strategies for 3D Printing

| **Feature / System** | **AN3DP – Active Nozzle Shape Control** | **Gradient Matters – Filament Diameter Adjustment** | **This work – Variable Diameter + Free Orientation** |
| --- | --- | --- | --- |
| **Method of diameter control** | Tendon-driven pins deforming flexible membrane → active diameter & shape change | Indirect control via print speed + nozzle height (no physical nozzle adjustment) | Gear-driven variable-diameter system; controlled directly without major G-code modifications |
| **Shape control** | Yes (cross-sectional shape and diameter both adjustable) | No (only deposition thickness varies) | **Diameter control + orientation freedom**. Shape control is possible with **one nozzle model** among the three interchangeable designs |
| **Orientation control** | Fixed nozzle axis; planar printing | Fixed nozzle axis; planar printing | Swivel-like extruder head allows **free-degree orientation** (printing on non-planar, curved, organ-like surfaces) |
| **Support structure need** | Reduced (thanks to contour fidelity) | **High** – requires added support layers to maintain fidelity | **Minimal** – orientation control reduces need for supports |
| **Mechanical complexity** | High (8 tendon actuators, membrane system, bulky) | Low (software/pathway-based, but limited) | **Moderate–Low** (gear system is compact, robust, stainless-steel compatible, easily heated for thermoplastics and bioinks) |
| **Adaptability to bioinks & heated materials** | Limited demonstration (mainly shear-thinning inks) | Not optimized for bioinks; mainly structural gradients | **Highly versatile**: compatible with soft bioinks and heated polymers due to stainless-steel body and heating capacity |
| **Resolution vs. speed tradeoff** | Improved via shape/dia. change, but mechanically complex | Limited; indirect diameter control constrains resolution–speed balance | Balanced; **real-time gear-driven dia. tuning** allows both high resolution and faster deposition |
| **Nozzle flexibility** | Single integrated nozzle, not interchangeable | Standard fixed nozzle | **Three interchangeable nozzle models** with different diameter-variation mechanisms; easy swap by unscrewing bottom |
| **Applications highlighted** | Multiscale engineering, gradient density structures, bone-like implants | Gradient structures, metastructures, flexible electronics, tissue-mimicking scaffolds | **Regenerative medicine + materials printing** – tissues, organ models, heated polymers, multifunctional fabrication |
| **Key disadvantages** | Bulky, complex actuation; not bioink-focused; no orientation freedom | Indirect, slow adaptation; support-dependent; planar only | Still needs further optimization for large-scale reliability, but offers the **widest versatility and ease of integration** |
